## Supplementary material for "Aberrant macrophage activation and maladaptive lung repair promote tuberculosis progression uniquely in the lung": Suppl. Figures

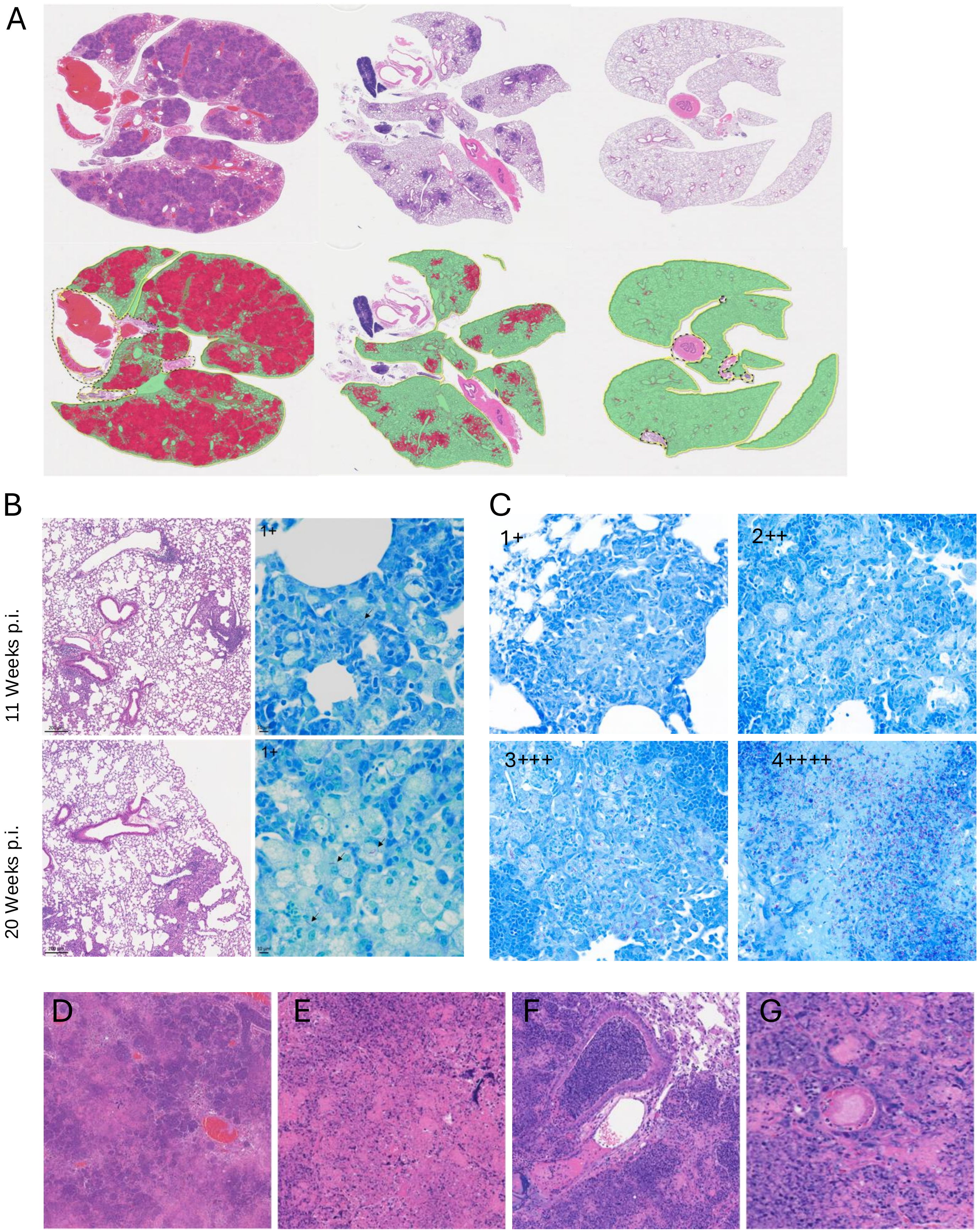


Supplemental Figure 1

**Supplemental Fig.1. Quantification of areas of inflammation and Mtb bacterial loads**

**A**. Representation for lesion quantification by digital classifier. Representative whole slide images of lungs of individual animals with severe pneumonia, moderate pneumonia, and lung that is within normal limits (Top panel, left to right, respectively). The bottom panel shows corresponding toggled images with areas classified as consolidation (pneumonia) by HALO image analysis software labeled in red. Note the absence of red labeling in the lung that is within normal limits (the right panel images), indicating the specificity of digital classification.

**B**. Representative histopathology and AFB load of infected C57BL/6J mice at 11 and 20 weeks post infection. Left two columns: H&E stain, 100X. Right two columns: Ziehl-Neelsen acid fast stain, 400X.

**C.** Representation of criteria of semi quantification of Mtb load in the lungs. Left upper, right upper, left lower, and right lower image represent Mtb loads of +, ++, +++ , and ++++, respectively. Please refer to the material and methods for the detailed description of the criteria.

**D - E.** Areas of complete tissue necrosis and the basophilic amorphous material adjacent to the thrombosed medium-caliber vessel (dystrophic mineralization), H&E.

**F.** The bronchioles are often occluded with large numbers of cell debris containing neutrophils and macrophages overflowed from the consolidated alveolar spaces. Occasional thrombi are observed in the lesions as shown in the center and 7 o'clock of the image, H&E.

**G.** Fibrinoid necrosis of a vessel in the area of necrosis. H&E.

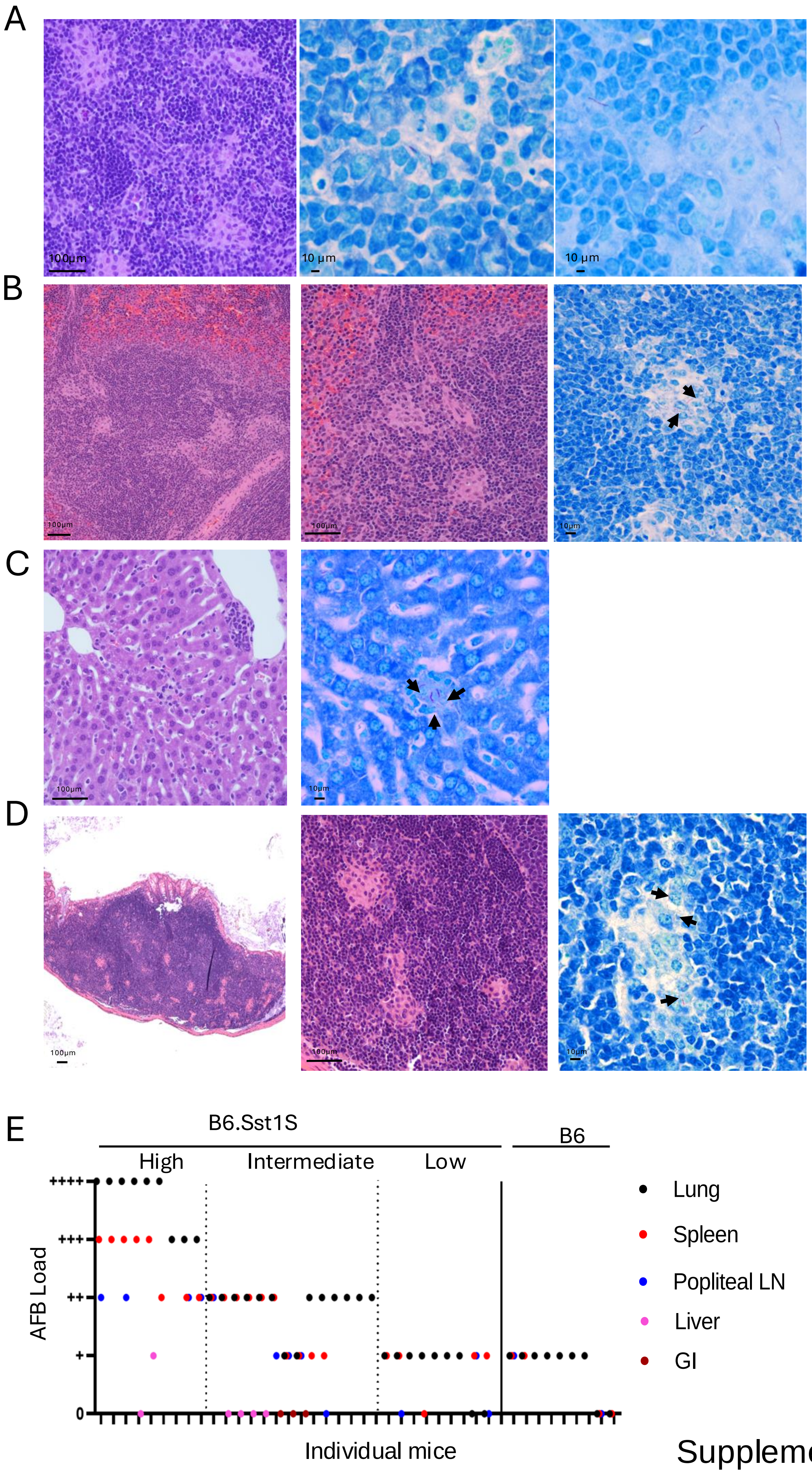


Supplemental Figure 2

**Supplemental Fig.2. Systemic dissemination of Mtb**

**A.** Representative histopathology and AFB load in popliteal lymph node of B6.Sst1S mouse at 6 weeks post hock infection with $10^6$ CFU of Mtb Erdman
Left panel - clusters of macrophages (microgranulomas) scattered in the peri-trabecular and medullary sinuses of popliteal lymph node (H&E stain, 100X). Middle – microgranuloma (H&E stain, 400X). Right panel - Ziehl-Neelsen acid fast stain, 400X. A few acid-fast bacilli (arrowhead) are scattered in the clusters of macrophages.

**B.** Representative images of histopathology and acid-fast bacilli load in the spleen of M. tuberculosis infected B6.Sst1S mice. Left image: clusters of macrophages (microgranulomas) scattered in the white pulp. H&E stain, 100X. Middle image: higher magnification of microgranulomas in the white pulp (H&E stain, 200X). Right image: A few single acid-fast bacilli (arrow) are scattered in the clusters of macrophages in the white pulp. Ziehl-Neelsen acid fast stain, 400X.

**C.** Representative images of histopathology and acid-fast bacilli load in the liver of M. tuberculosis infected B6.Sst1S mice. Left image: a rare cluster of macrophages in the sinusoids. H&E stain, 200X. Right image: A few single acid-fast bacilli (arrow) are scattered in the clusters of macrophages in the sinusoids.

**D.** Representative images of histopathology and acid-fast bacilli load in the cecum of M. tuberculosis infected mouse. Left image: multifocal small clusters of macrophages (microgranulomas) scattered in the GALT (gut-associated lymphoid tissues). H&E stain, 100X. Middle image: higher magnification of microgranulomas in the GALT (H&E stain, 200X). Right image: A few single acid-fast bacilli (arrow) are scattered in the clusters of macrophages in the GALT. Ziehl-Neelsen acid fast stain, 400X.

**E**. Histopathological scores indicating Mtb loads across different organs in B6 (n=9) and B6.Sst1S (n=32) mice. Each mark on the X-axis represents an individual mouse.

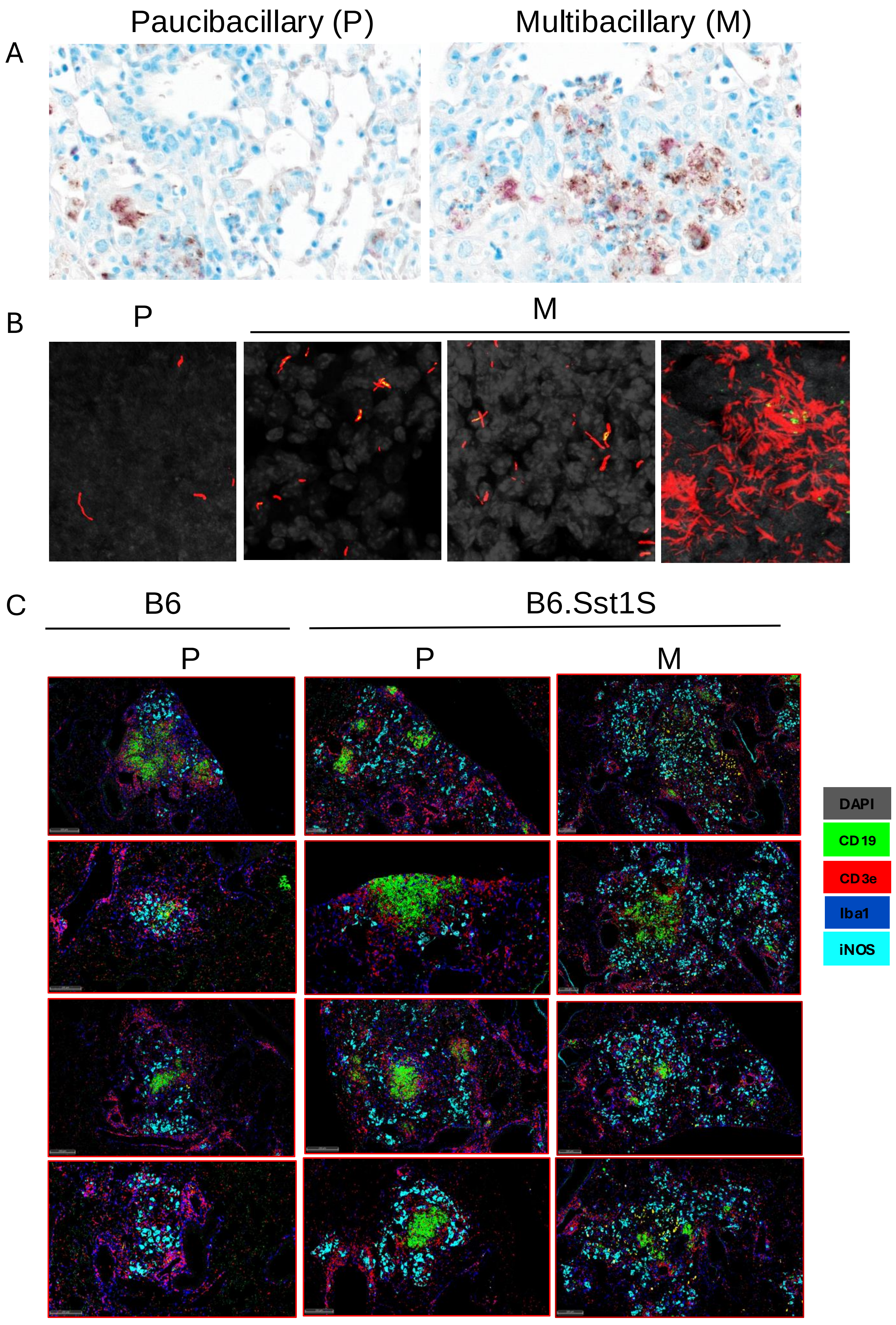


Supplemental Figure 3

**Supplemental Fig.3. Comparative Analysis of Paucibacillary and Multibacillary TB Lesions in B6 and B6.Sst1S Mice**

**A.** Duplex immunohistochemistry (Mtb polyclonal) combined with Acid Fast Bacilli (AFB) histochemical stain. Paucibacillary (P) and multi-bacillary (M) lesions. Mtb antigen and AFB colocalized within identical anatomical compartments (i.e., cytoplasm of macrophages). Images clearly show that Mtb bacilli and antigen loads were low in C lesions, significantly increasing in NC lesions. All images taken at 400X magnification.

**B.** 3D confocal images of 50 μm thick cleared lung sections of paucibacillary and multi-bacillary intermediate and terminal TB lesions of mice infected with Mtb reporter strain expressing replication marker (SSB-GFP, *smyc'*::mCherry). Replication reporter (SSB-GFP) is shown as green dots; Mtb (smyc':: mcherry) is red and nuclei in grayscale (DAPI). The fraction of the replication reporter-positive Mtb in intermediate M lesions increased to 30 - 50% (left and middle panels). Scale bar- 50μm

**C**. fmIHC multiple images for wild type B6, B6.Sst1S- paucibacillary (P), and B6.Sst1S- multi-bacillary (M) TB lesions at 200X magnification showing accumulation of activated macrophages (iNOS) in B6.Sst1S- (M), with dissolution of discrete lymphoid follicles. DAPI-Gray; CD19 - green; CD3 epsilon (CD3ε) - red; inducible nitric oxide (iNOS) -teal; ionized calcium-binding adaptor molecule 1 (Iba1) - blue. All images taken at 10X magnification.

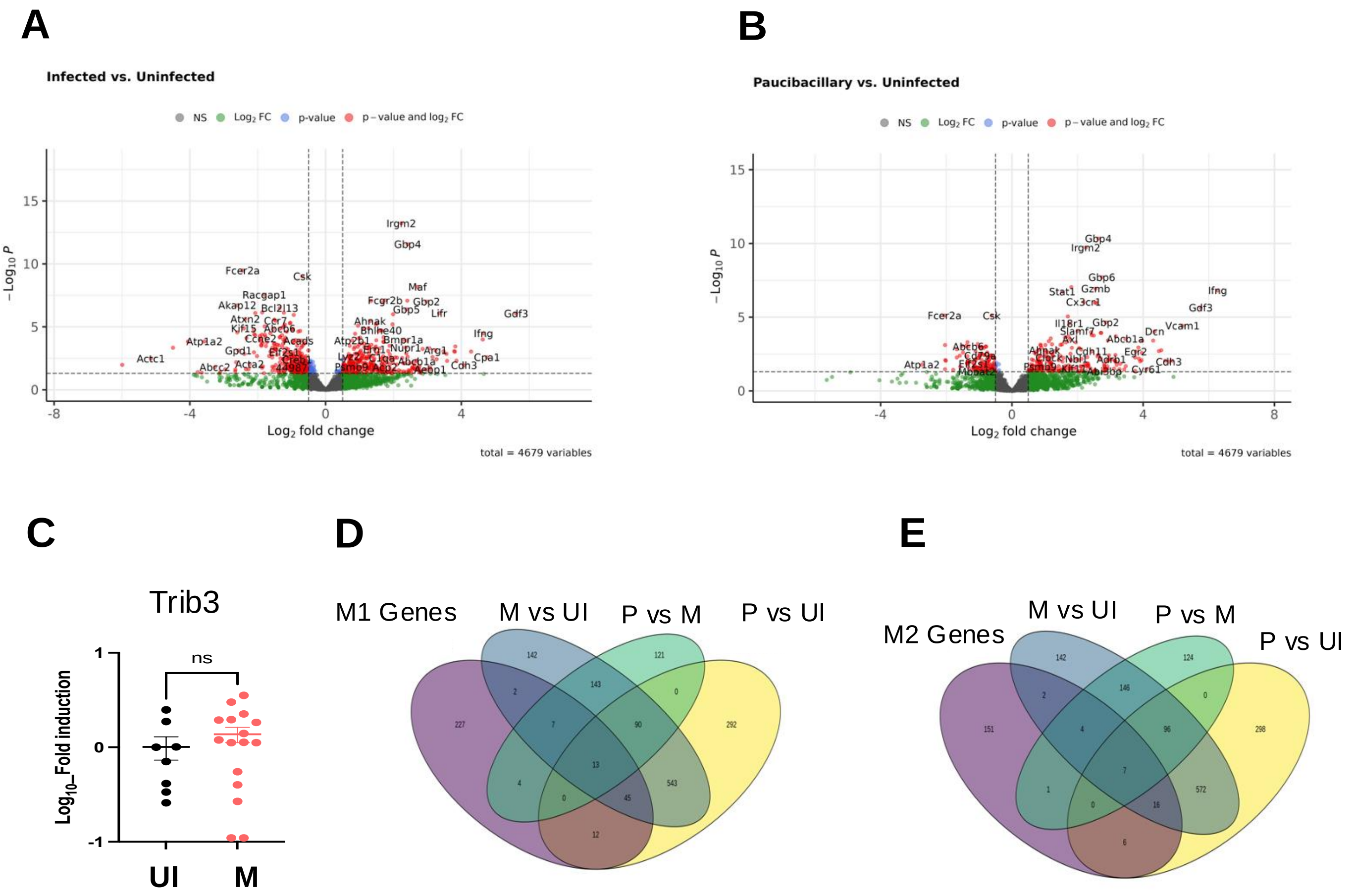


Supplemental Figure 4

**Supplemental Figure 4. Blood transcriptome analysis of circulating myeloid cells**

**A - B.** Volcano plots showing differentially expressed genes in blood samples from M. tuberculosis-infected mice compared to non-infected controls (A) and blood samples from M. tuberculosis-infected mice with paucibacillary lung lesions compared to non-infected controls (B). Each point represents a single gene, with the x-axis indicating log2 fold change and the y-axis showing -log10(p-value). Significantly downregulated and upregulated genes are highlighted in red (left and right, respectively), while non-significant genes are shown in gray. Significance thresholds are set at $p < 0.05$ and fold change > ±1.5.

**C.** Expression of Trib3 mRNA in blood of noninfected control and Mtb infected mice with multi-bacillary stage lung lesions.

**D - E.** Quantification of M1 and M2 macrophage-associated genes, as determined by blood transcriptomics, in mice with paucibacillary and multibacillary TB lung lesions compared to non-infected controls, showing overlap and distinctions in gene expression associated with each macrophage polarization state.

The statistical significance was performed by two-tailed unpaired t test (Panel C). Significant differences are indicated with asterisks (*, $P < 0.05$; **, $P < 0.01$; ***, $P < 0.001$; ****, $P < 0.0001$).

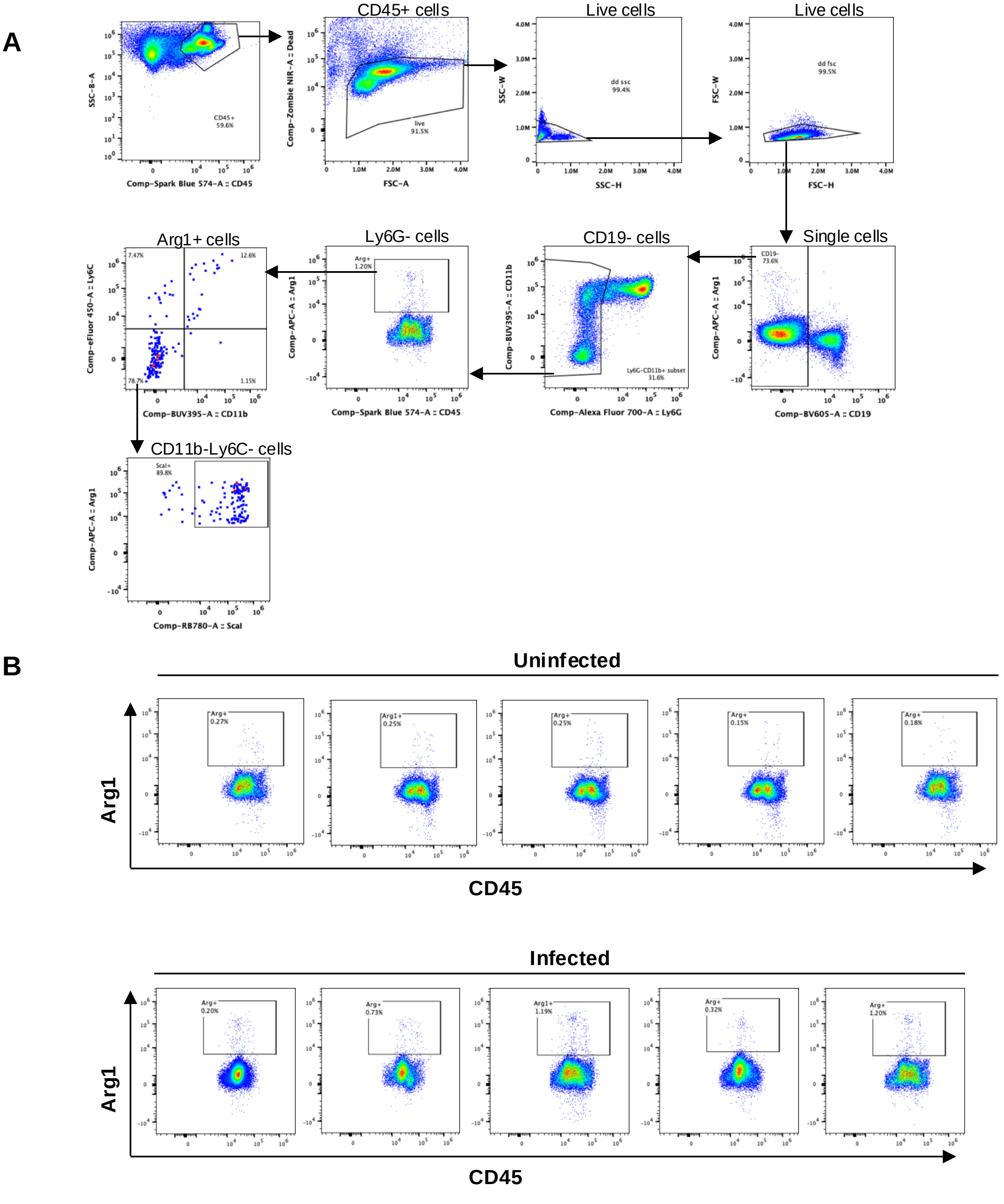


Supplemental Figure 5

**Supplemental Fig.5. Mtb infection leads to increase in Arg1 expressing cells in bone marrow.**

**A.** Gating strategy for flow cytometry data analysis.

**B.** The CD45 vs Arg1 plot showing the increase in Arg1 expressing cells in bone marrow upon Mtb infection, five mice of each group.

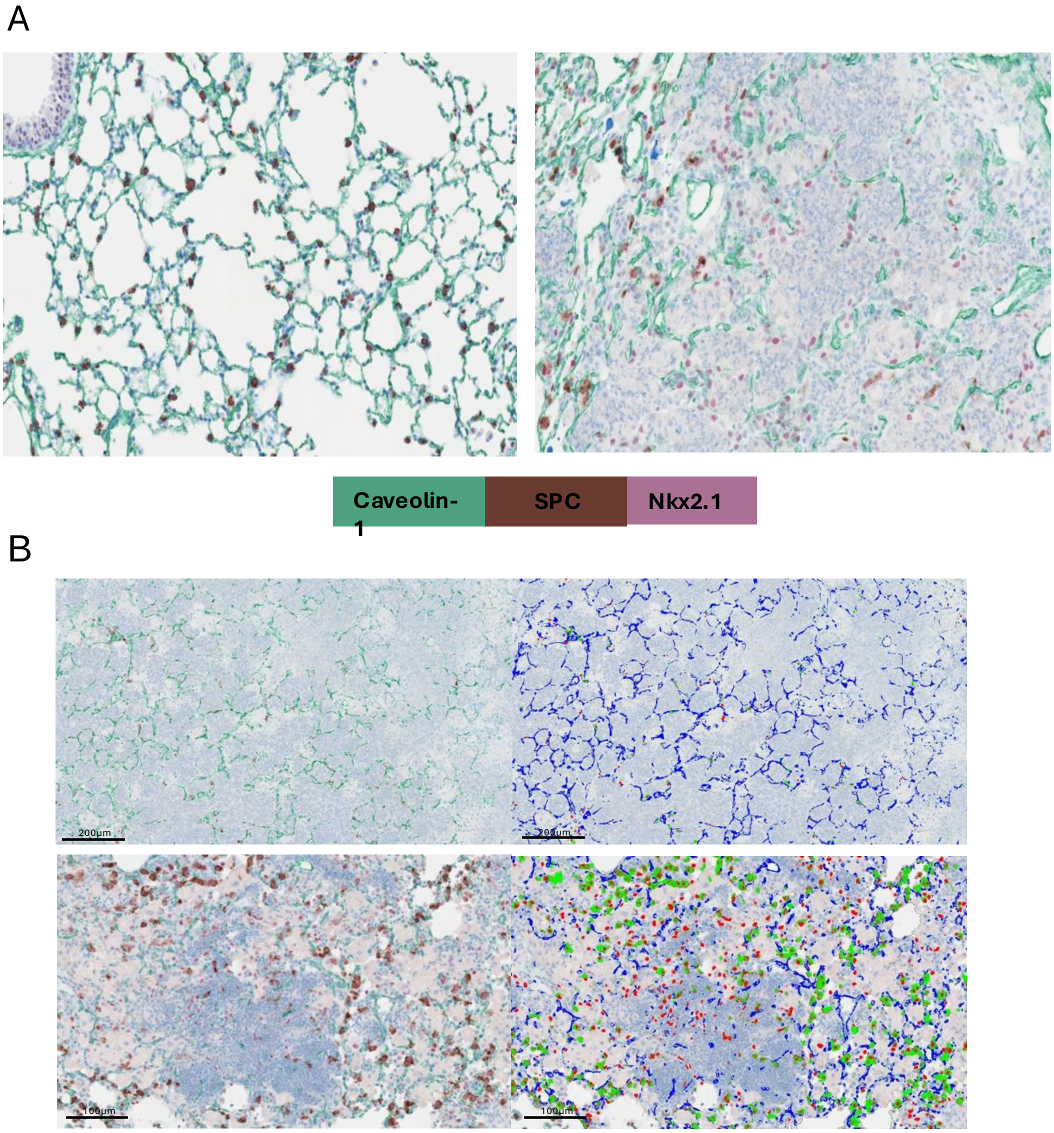


Supplemental Figure 6

**Supplemental Fig.6. Dysplastic lung epithelial cells in PTB lesions of B6.Sst1S mice**.

**A.** Comparison of representative images of chromogenic immunohistochemistry staining of Nkx2.1 (magenta), SPC (brown) and Caveolin-1 (green) in unaffected pulmonary parenchyma in a B6 mouse versus granulomatous pneumonia lesions in a B6.Sst1S,ifnβ-YFP mouse.

**B.** Representative images of chromogenic immunohistochemistry staining of Nkx2.1 (magenta), SPC (brown) and Caveolin-1 (green) of necrosuppurative pneumonia of Mtb infection in a B6.Sst1S,ifnβ-YFP mouse (top panel) and granulomatous pneumonia in a B6.Sst1S mouse (bottom panel). Left panel-top: 100X; bottom: 200X. Right panel-Corresponding toggled digital classification of the images in the left panel showing indicative IHC signals recognized by the trained algorithm (Nkx2.1 recognized in red, SPC recognized in green and Caveolin-1 recognized in blue).

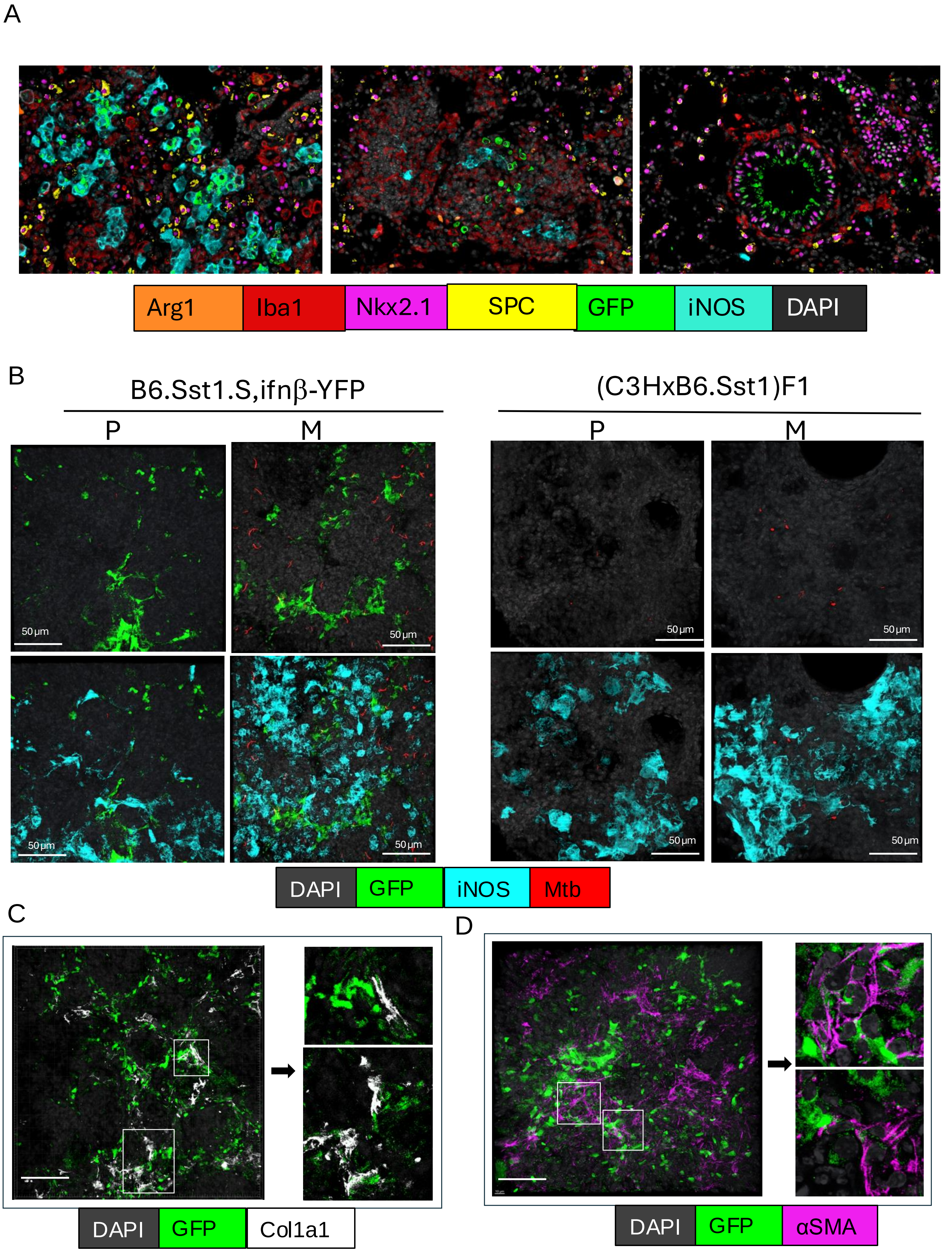


Supplemental Figure 7

**Supplemental Fig.7. The expression of the IFNβ reporter YFP in TB lesions. Majority of YFP/IFNβ+ cells are iNOS+ activated macrophages.**

**A.** Additional cell populations illustrated to express GFP/IFN reporter (iNOS+ macrophages-left; perivascular Iba- mononuclear cells –middle; airway epithelium-right).

**B.** 3D confocal images of cleared 50 μm thick lung sections of B6.Sst1S,ifnβ-YFP and (C3H x B6.Sst1) F1 mice. Endogenous YFP (seen only in B6.Sst1.S,ifnβ-YFP) is shown in green, iNOS staining in teal blue, reporter bacteria in red (smyc':: mcherry) and nuclei in grayscale (DAPI). Scale bar- 50µm.

**C**. 3D confocal image of cleared 50 μm thick lung sections of B6.Sst1S,ifnβ-YFP mice stained with Col1a1 antibody; on the right – individual images of Col1a1+ cells adjacent to YFP (denoted by white boxes on the left panel). Scale bar- 50µm.

**D**. 3D confocal images of cleared 50 μm thick lung sections of B6.Sst1S,ifnβ-YFP mice stained with myofibroblast marker αSMA antibody. Endogenous YFP is shown in green, αSMA staining in magenta, and nuclei in grayscale (DAPI); on the right – individual images show that αSMA+ cells are adjacent to YFP (denoted by white boxes on the left panel). Scale bar- 50µm.

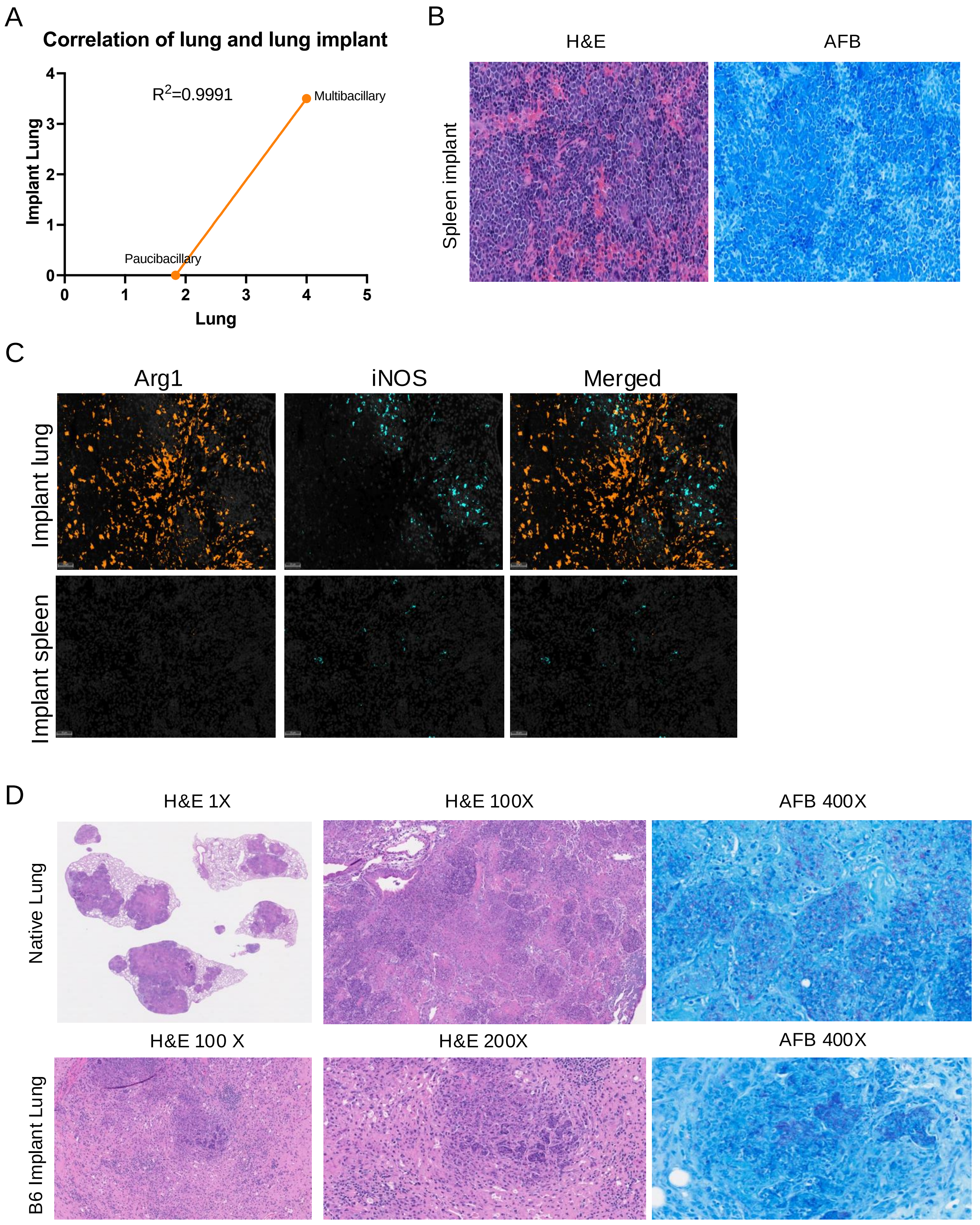


Supplemental Figure 8

**Supplemental Fig.8. Characterization of TB lesions in lung and spleen implants**

**A.** Correlation of acid-fast bacilli (AFB) score in lungs and respective lung implants of mtb-infected mice. AFB score were annotated via Ziehl-Neelsen staining and microscopic observations in both primary lung tissues and implants.

**B.** Representative hematoxylin and eosin (H&E) and acid-fast bacilli (AFB) staining of spleen implant. 200x total magnification.

**C**. fmIHC images of Arg1+ and iNOS+ cells in lung and spleen implants of Mtb infected mice. Arg1+ cells - orange color and iNOS+ - teal color.

**D**. Corresponding TB lesions in the native lung of B6.Sst1S recipient mice (upper panel) and in implanted lung tissue isolated from B6 donors (lower panel). Implanted B6 lung tissue developed the necrotic lesions containing numerous Mtb. H&E and AFB staining.
