## Supplementary material for "Aberrant macrophage activation and maladaptive lung repair promote tuberculosis progression uniquely in the lung": Suppl. Tables

**Supplemental Table 1. Histopathology and acid-fast bacilli load in lung of all animals.**

| **Time post infection** | **Genotypes** | **WNL (AFB^2^ in lung)** | **mononuclear infiltrates^1^ (AFB^2^ in lung)** | **Granulomatous pneumonia only (AFB^2^ in lung)** | **Necrosuppurative pneumonia**  **(AFB^2^ in lung)** | **AFB^2^ lung** | | | | |
| --- | --- | --- | --- | --- | --- | --- | --- | --- | --- | --- |
| 11 weeks | C57BL/6J | 1/4 (-, 1/1) | 1/4 (-, 1/1) | 2/4 minimal to moderate (-, 1/2; +, 1/2) | 0/4 | 3/4 | 1/4 | 0/4 | 0/4 | 0/4 |
|  | B6.Sst1.S and B6.Sst1.S,ifnb-YFP | 0/14 | 1/14 (-, 1/1) | 10/14 mild to marked^3^ (-, 1/10; +, 3/10; ++, 3/10, +++, 2/10; ++++ 1/10) | 3/14 (++++, 3/3) | 2/14 | 3/14 | 3/14 | 2/14 | 4/14 |
| 20 weeks^4^ | C57BL/6J | 0/5 | 0/5 | 5/5 mild to moderate (+, 5/5) | 0/5 | 0/5 | 5/5 | 0/5 | 0/5 | 0/5 |
|  | B6.Sst1.S and B6.Sst1.S,ifnb-YFP | 0/15 | 0/15 | 13/15 minimal to marked^3^ (+, 4/13; ++, 7/13; +++, 2/13) | 2/15 (++++, 2/2) | 0/15 | 4/15 | 7/15 | 2/15 | 2/15 |

^1^ peribronchiolar, perivascular and/or interstitial

^2^ AFB: acid-fast bacilli. See material and method for semi-quantification criteria.

^3^ with occasional necrosis, occasional to frequent cholesterol cleft and neutrophil influx

^4^ The animals in the 20 week group have a range of time post inoculation from 19 weeks, 20 weeks, 24 weeks and 26 weeks.

C57BL/6J infected animals developed granulomatous pneumonia that affected 0.59% to 5.58% of total examined pulmonary parenchyma, with no detectable to low (- to +) Mtb load. The Mtb load in B6.Sst1.S mice was increased compared to C57BL/6J mice, with the majority ranging from ++ to ++++ (n=20). Compared to the ++++ Mtb load in the animals that developed necrosuppurative pneumonia, B6.Sst1S mice with granulomatous pneumonia displayed a maximum +++ Mtb load, with one exception in an animal that developed marked granulomatous pneumonia, which displayed a ++++ Mtb load (**Suppl.Table 3**). Rarely, B6.Sst1.S mice had no detectable Mtb in the examined sections (n=2); and a few animals had a low (+) Mtb load (n=7).

**Supplemental Table 2. Histopathology and acid-fast bacilli load in spleen of examined animals.**

| **Time post infection** | **Genotypes** | **WNL**  **(AFB^2^ in spleen)** | **Microgranulomas in the white pulps (AFB^2^ in spleen)** | **AFB^2^ spleen** | | | |
| --- | --- | --- | --- | --- | --- | --- | --- |
| 11 weeks | C57BL/6J | 0/4 | 4/4 rare to small numbers  (-, 2/4; +, 2/4) | 2/4 | 2/4 | 0/4 | 0/4 |
|  | B6.Sst1.S and B6.Sst1.S,ifnb-YFP | 4/14 (-, 1/4; +, 3/4) | 10/14 rare to medium numbers (+, 3/14; ++, 4/14, +++, 3/14) | 1/14 | 6/14 | 4/14 | 3/14 |
| 20 weeks | B6.Sst1.S and B6.Sst1.S,ifnb-YFP | 0/6 | 6/6 medium to large numbers (++, 4/6; +++, 2/6) | 0/6 | 0/6 | 4/6 | 2/6 |

Three to four sections of spleen from each C57BL/6J infected mice (n=4 at 11 week-post infection) and B6.Sst1.S infected mice (n=14 at 11-wpi; and n=6 at 20 week-post infection) were examined. The most consistent observation across all infected animals in animals with concurrent pulmonary lesions was the presence of rare to medium numbers of microgranulomas characterized by nodular clusters of macrophage aggregates multifocally within the white pulp (**Suppl. Fig. 2A**). Most of the spleen microgranulomas contained low to intermediate Mtb load (+ to ++) with single individualized acid-fast bacilli (**Suppl. Fig. 2A**). Five B6.Sst1.S infected mice animals that developed necrosuppurative pneumonia with high Mtb load in the lung (++++) had also high Mtb load (+++) in the spleen, likely reflecting high Mtb load in the systemic circulation (**Fig. 2D; Suppl. Table 4**). Four B6.Sst1.S infected mice had no significant histopathologic findings in the spleen. In summary, in B6.Sst1.S infected mice, there were 13 out of 14 animals at 11 wpi and 6 out of 6 animals at 20 wpi with detectable Mtb in the spleen. Two of four examined C57BL/6J infected mice had no detectable Mtb in the spleen, and the other two mice had rare individualized (+) Mtb.

**Supplemental Table 3. Histopathology and acid-fast bacilli load in popliteal lymph nodes of examined animals.**

| **Time post infection** | **Genotypes** | **WNL**  **(AFB^2^ in popliteal lymph node)** | **Sinus histiocytosis to microgranulomas (AFB^2^ in popliteal lymph node)** | **AFB^2^ popliteal lymph node** | | | |
| --- | --- | --- | --- | --- | --- | --- | --- |
| 11 weeks | C57BL/6J | 0/2 | 2/2 mild to moderate (-, 1/2; +, 1/2) | 1/2 | 1/2 | 0/2 | 0/2 |
|  | B6.Sst1.S and B6.Sst1.S,ifnb-YFP | 1/9 (-, 1/1) | 8/9 mild to marked (-, 2/8; +, 1/8; ++, 5/8) | 3/9 | 1/9 | 5/9 | 0/9 |
| 20 weeks | B6.Sst1.S and B6.Sst1.S,ifnb-YFP | 0/6 | 6/6 medium to large numbers (++, 4/6; +++, 2/6) | 0/6 | 0/6 | 4/6 | 2/6 |

In the popliteal lymph nodes, at 11 week post infection, there was mild to moderate sinus histiocytosis, with sporadic microgranulomas in the peritrabecular and medullary sinuses. Among nine B6.Sst1.S infected animals with popliteal lymph nodes available for examination, six (66.7%) contained rare to low numbers of single, acid-fast bacilli (+ to ++) in the microgranulomas; and in two infected C57BL/6J mice, one had no detectable Mtb in the popliteal lymph node; while the other one had rare (+) individual acid-fast bacilli.

**Supplemental Table 4: Excel file.**

**Differentially expressed genes (DEGs) in total ROIs in Intermediate vs Advanced TB lesions.**

**Supplemental Table 5 Pathways upregulated in Advanced vs Intermediate TB lesions (Total ROIs).**

MSigDB Hallmark 2020

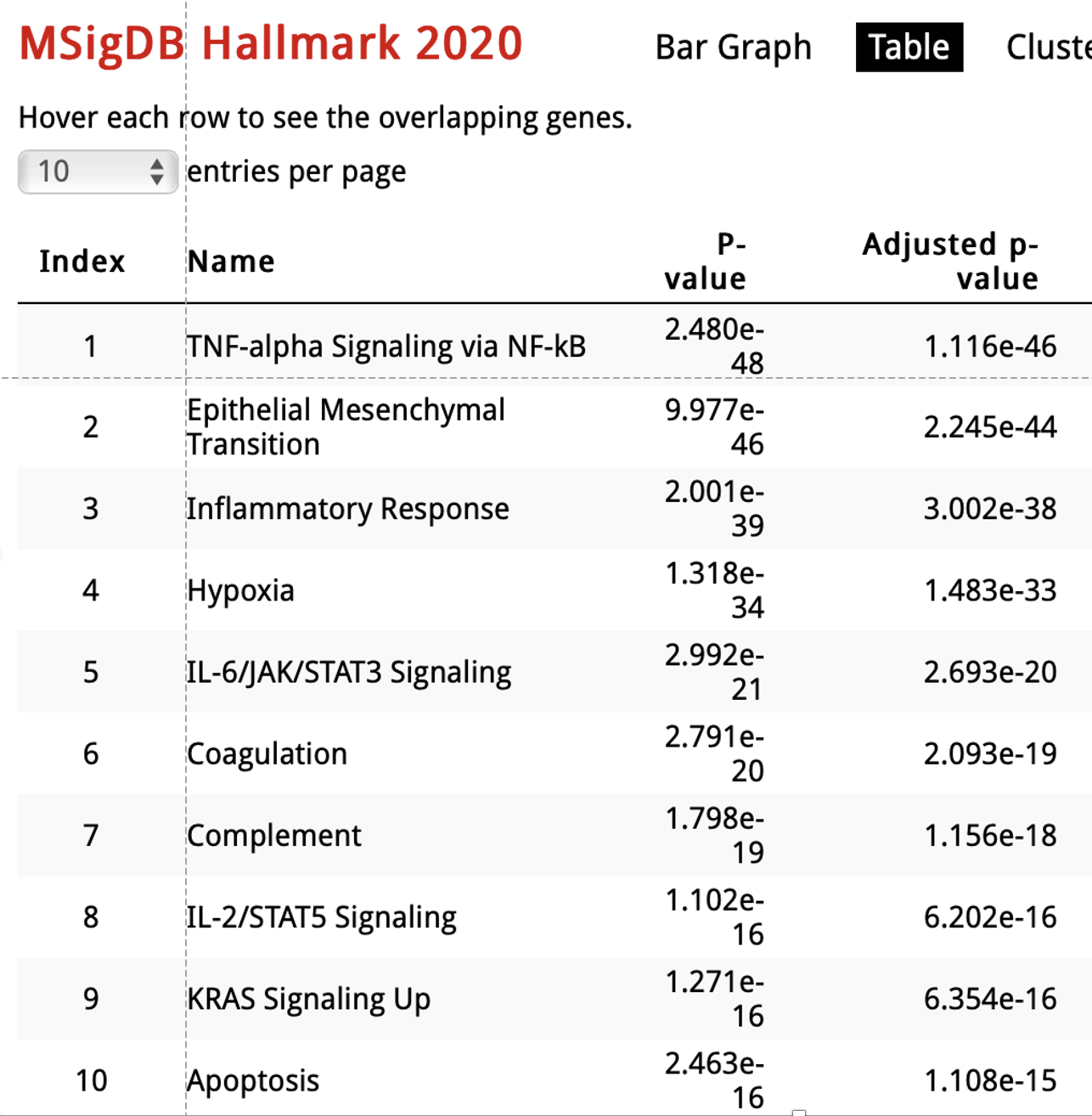

**Supplemental Table 6. The list of antibodies used for staining cells for flow cytometry.**

| **Antibody Name** | **Source: Catalog number** |
| --- | --- |
| Anti-CD11b BUV395 | BD Biosciences: 565976 |
| Anti-Ly6C eFluor 450 | Thermo Scientific: 50-246-021 |
| Anti-Sca-1 RB780 | BD Biosciences: 569230 |
| Anti-MHC II BV650 | Thermo Scientific: 41-653-2180 |
| Anti-CD115 NovaFluor Yellow 730 | Thermo Scientific: M041T02Y07-A |
| Anti-CCR2 PE-Fire 640 | Biolegend: 150639 |
| Anti-CD206 PE-Fire 700 | Biolegend: 141741 |
| Anti-Lifr PE-Cy7 | Biolegend: 158903 |
| Anti-Ly6G Alexa Fluor 700 | Biolegend: 127621 |
| Anti-CD11c BV785 | Biolegend: 117335 |
| Anti-CX3CR1 APC-Fire 810 | Biolegend: 149053 |
| Anti-CD19 BV605 | Biolegend: 115539 |
| Anti-CD45 Spark Blue 574 | Biolegend: 103183 |
| Anti- Arg1 APC | Thermo Scientific: 17-369-780 |

**Supplemental Table 7. Cell counts of Arg1 expressing cells in bone marrow**

|  | **CD45+Arg+** | **CD11b+Ly6C+** | **CD11b-Ly6C-Sca1+** |
| --- | --- | --- | --- |
| **Uninfected mice** | 59 | 8 | 26 |
|  | 54 | 7 | 27 |
|  | 35 | 1 | 19 |
|  | 45 | 4 | 21 |
|  | 27 | 6 | 13 |
| Average | 44 | 5.2 | 21.2 |
| **Infected mice** | 160 | 45 | 62 |
|  | 174 | 22 | 123 |
|  | 119 | 65 | 33 |
|  | 282 | 44 | 174 |
|  | 100 | 25 | 57 |
| Average | 167 | 40.2 | 89.8 |
